# Exclusive liquid repellency isolation (ELRi) microliter scale isolation technique unmasks T-cell migration defects to CCL21 in pediatric asthma

**DOI:** 10.64898/2026.05.26.727898

**Authors:** Terry D Juang, Brendan M. Joyce, Neha Prasad, Yuanshan Li, Fauzan Ahmed, Gina M Crisafi, Christine M Seroogy, James E Gern, Irene M Ong, David J Beebe, Sheena C Kerr

## Abstract

Exclusive Liquid Repellency Isolation (ELRi) is a novel, chip-free platform designed for efficient microscale immune cell isolation. Leveraging the inherent exclusive liquid repellency (ELR) properties of polypropylene tubes with an oil overlay, ELRi ensures that small sample volumes, as low as 8 μL, have no contact with container walls, preventing cell loss. Here we describe ELRi and demonstrate magnetic bead-based isolation of diverse immune cells, including T cells, and monocytes, from whole blood. The platform’s utility is highlighted in its application to pediatric asthma research, where sample volume is highly restricted. It has been reported that the T cell homing receptor CCR7 is downregulated in cells from asthmatic patients, but a direct functional link to impaired cell migration remains unconfirmed (1, 2). Using ELRi-isolated cells, we provide the first functional evidence that T cells from asthmatic children exhibit significantly impaired chemotaxis toward the CCR7 ligand, CCL21. By simplifying RBC depletion and enabling such functional assays alongside RNA sequencing from the same tiny sample, ELRi overcomes the limitations of large-volume flow cytometry or cell-specific sorting methods. Easily integrated into lab workflows and scalable for various needs, ELRi facilitates more frequent, minimally invasive, and functionally informative immune profiling of restricted volume samples.

## Introduction

Despite the growing demand for frequent, minimally invasive blood analyses, robust methods for obtaining immune populations from microliter-scale samples remain limited. One of the main challenges in microscale immunology is the overwhelming presence of red blood cells (RBCs), which outnumber white blood cells (WBCs) by roughly 2,000:1 (3). RBC contamination can complicate downstream analyses, including imaging, immunophenotyping, and transcriptomics. Traditional protocols for immune cell isolation such as density gradients require milliliter volumes and are suboptimal when translated into microliter volumes. This results in cell losses during the multiple wash steps and compromised purity of the samples as surface tension dominates fluid behavior preventing reliable fractionation (4). Consequently, microfluidics emerges as a potential solution, enabling stable control over ultra-small volumes and facilitating the targeted isolation of specific cell types. Various microfluidic devices have been developed to address small-volume separations, often leveraging cell size, deformability, or other biophysical properties (5, 6). While these approaches have successfully isolated neutrophils directly from a drop of blood and yielded valuable insights into their behavior (e.g., chemotaxis, neutrophil extracellular trap (NET) formation) (7–13), their reliance on specific cell properties such as the high deformability of neutrophils restricts the isolation of other immune subsets i.e T cells, or monocytes. While T cells have been isolated in a microfluidic platform from blood of acute lymphoblastic lymphoma patients, T cell separation was based on the size of T cells relative to enlarged lymphoblasts, therefore would not be applicable to non-leukemic samples (14). Magnetic bead-based isolation strategies provide broader applicability by relying on immunoaffinity rather than cell-specific biophysical traits. However, many magnetophoretic approaches require larger volumes. These limitations create a need for a platform that can handle microliter-scale samples, remove RBCs, and isolate multiple immune cell subsets rapidly and reliably, all without extensive sample preparation or specialized instrumentation.

These constraints are especially important for immunological studies, particularly in pediatric or neonatal contexts where amounts of blood that can be safely drawn are restricted (15). The ability to use residual small volumes of blood from clinical tests would open potential to study immune changes temporally across a disease course such as patients with sepsis in the ICU who undergo frequent blood sampling for clinical testing where residual volumes are insufficient for traditional analyses. Obtaining robust immunological data from a few microliters of blood can enable frequent, minimally invasive sampling (16–18) that captures dynamic immune states over time. Achieving this requires methods that efficiently isolate specific immune cell populations from low-volume blood samples without compromising cell function or downstream analytical quality.

Exclusive Liquid Repellency (ELR) (19) describes the condition in which an aqueous droplet is completely repelled from a solid surface by an immiscible continuous oil phase. Thus, ELR provides a stable environment that prevents evaporation and biofouling, enabling the isolation and manipulation of microscale samples. Here, we leverage this to create exclusive liquid repellency isolation (ELRi) permitting direct magnetic bead isolation of immune cells within an aqueous droplet with a volume as low as 8 μL in approximately 5 minutes. Further, ELRi can be applied to multiple immune cell types including T cells and monocytes. Compatibility of ELRi with existing microscale analysis platforms, such as the kit on a lid assay (KOALA) (8, 12) permits functional studies to be performed with these microscale immune isolations. In addition to functional assays, RBC-depleted and immune cell-enriched samples obtained through ELRi are highly compatible with RNA sequencing (RNA-seq). Sequencing whole blood samples can result in globin transcripts comprising 70–90% of the data (20, 21) and existing approaches to deplete globin RNA from cDNA libraries (22, 23) are not compatible with ultra-low input RNA-seq library preparation kits. ELRi can effectively deplete RBCs from microliter-volume blood samples before RNA sequencing.

In this study, we introduce ELRi, as a rapid, one-step workflow for isolating immune-cell subsets or depleting RBCs from ultra-low-volume blood samples. We demonstrate how the approach preserves cellular functionality and supports immediate downstream assays. We used ELRi to isolate T cells from a pediatric cohort of asthmatic and non-asthmatic children for downstream functional assays. We observed a previously unidentified T cell migration defect in asthmatic children, providing new insights into pediatric T cell biology in asthma. Thus, ELRi as a practical platform for microscale immune profiling can facilitate investigation of immune cell biology in pediatric or other restricted volume patient cohorts.

## Materials and Methods

### Blood Collection

Human whole blood samples were collected from healthy adult donors under a protocol approved by the Institutional Review Board (IRB) at the University of Wisconsin-Madison (2020–1623). For pediatric blood collection, samples were collected from children between the ages of 4-16 years under a protocol approved by the IRB at the University of Wisconsin-Madison (2022–0307). Children were classified as having a history of asthma or allergy based on a caregiver-administered questionnaire. Informed consent was obtained from all participants. Blood was drawn into ACD-coated Vacutainer tubes (BD).

### Immune Cell Isolation Kits and Reagents

All magnetic negative selection kits—including EasySep™ Human T Cell Isolation Kit (Catalog #17951), EasySep™ Human Monocyte Isolation Kit (Catalog #19669), and EasySep™ RBC Depletion Kit (Catalog #18170) were purchased from StemCell Technologies. RapidSpheres™ magnetic particles, as well as antibody cocktails, were used following the manufacturer’s guidelines. PBS, RPMI, and ImmunoCult™ media were obtained from standard suppliers (Thermo Fisher).

### Exclusive Liquid Repellency Isolation (ELRi) Setup and Procedure

The ELRi method was established using standard 0.6 mL polypropylene Eppendorf tubes, with antibody cocktail and magnetic bead volumes proportionally adjusted according to manufacturer recommendations. Whole blood samples were incubated with antibody cocktail and RapidSpheres at ratios of 50 µL antibody cocktail and 100 µL beads per 1 mL of blood for 5 minutes at room temperature. For small-volume isolations, reaction volumes were proportionally scaled down, typically involving 10 µL blood combined with 0.5 µL antibody cocktail and 1 µL RapidSpheres. Subsequently, 10 µL phosphate-buffered saline (PBS) was added, yielding a final reaction volume of approximately 20 µL.

An 8 µL aliquot of this reaction mixture was gently pipetted into the bottom of a 0.6 mL Eppendorf tube prefilled with 50 µL silicone oil (5 cSt, Sigma-Aldrich), ensuring the aqueous droplet remained suspended at the oil interface without contacting the tube walls. Magnetic separation was performed using a custom-made “magnetic tip,” constructed by embedding a small magnet (KCJ Magnetics, D12-N52; diameter 1/16”) into a standard P200 pipette tip, sealed at the end with wax. Upon gentle contact of this magnetic tip with the droplet, magnetically labeled cells—including erythrocytes and other unwanted cells—rapidly clustered toward the tip within approximately 1 minute, while the unlabeled target cells remained at the bottom of the droplet.

Subsequent withdrawal of the magnetic tip upward caused the original 8 µL droplet to split into two distinct compartments within the oil phase: a smaller (∼2 µL) upper fraction containing magnetically labeled cells (primarily erythrocytes and other unwanted cells), and a larger (∼6 µL) lower fraction enriched with unlabeled target cells. The magnetic tip carrying the unwanted cells was removed and cleaned using disinfectant wipes (CaviWipes), enabling additional isolation rounds if needed. The ∼6 µL cleared droplet containing isolated target cells was directly suitable for downstream analytical assays such as flow cytometry, imaging, or RNA sequencing. For benchmark comparisons of ELRi with macroscale isolations, T cells were isolated from 10 mL of blood using the same negative isolation kits and following the manufacturers recommendations.

### Scaling up ELRi (ELRi XL)

When isolating less abundant cell populations (e.g., monocytes) or when greater cell numbers are required for downstream assays, the blood volume can be scaled up while still using minimal collection methods such as a fingerprick, heelstick, or Tasso+ (16–18). Maintaining the same 50 µL antibody cocktail and 100 µL RapidSpheres per 1 mL blood ratio, 40 µL of whole blood is incubated with 2 µL of antibody cocktail and 4 µL of RapidSpheres for 5 minutes at room temperature. Then, 40 µL of PBS is added, bringing the total reaction volume to ∼86 µL.

From this mixture, 70 µL is carefully transferred into a 5 mL Eppendorf tube prefilled with 200 µL silicone oil (5 cSt, Sigma-Aldrich). A larger custom-built magnet (KCJ Magnetics, D33-N52, 3/16” diameter) embedded in a P1000 pipette tip and sealed with wax is used for separation. Within about one minute, magnetically labeled, unwanted cells cluster toward the tip, leaving unlabeled target cells at the oil interface. If a volume greater than 70 µL is needed, multiple tubes can be processed back-to-back, providing flexibility while still keeping each isolation step rapid. The enriched droplet of unlabeled cells is then ready for downstream analyses such as flow cytometry, imaging, or RNA sequencing.

### Device Coating and KOALA Preparation

The KOALA platform (12)(Figure 5) was used to quantify migration of purified T cells. Devices were fabricated from polydimethylsiloxane (PDMS) as previously described (12, 24). With the PDMS device placed on an IBIDI µ-Slide (Cat. #82101) and kept on ice, channels dedicated to T cell imaging were coated with 6 µL recombinant human ICAM-1 (10 µg mL⁻¹ in ddH₂O, RCD Systems). Devices were stored overnight at 4°C in a refrigerator. The next day, coating solutions were aspirated and ∼4 µL of PBS was introduced into each channel. T cells were isolated from whole blood using the ELRi workflow and labeled with Hoechst (1:1000). Cells were than loaded into each channel by adding 1 μL aliquots every minute, totaling 8 μL per channel. After a 10-min rest at room temperature, devices were transferred to the microscope for time-lapse imaging.

### Migration Assay and Imaging

To establish chemotactic gradients, 7.5 µL of neutralized type I collagen (Corning 354249, 10 mg mL⁻¹ stock mixed 1 : 1 with HEPES-buffered PBS to pH ≈ 7.2, yielding 4 mg mL⁻¹ final) supplemented just before casting with 5 nM CCL21 (from PBS stock) for T cell assays was dispensed onto each KOALA lid, whose wells contain only chemoattractant. PBS droplets were placed around the lid to reduce evaporation. Collagen polymerization occurred at room temperature for 30–60 minutes. The KOALA µ-Slide was then transferred to a Nikon TiE1 microscope with a heated, humidified, and O_2_-controlled stage set to 37°C. Time-lapse imaging (every 2 minutes for up to 3 hours) was performed using a 20X objective in brightfield and blue fluorescence channels. Cell trajectories were analyzed using FIJI and TrackMate, with linking/gap distances set at 20 μm and the IBIDI Chemotaxis and Migration Tool for Center of Mass (COM) values. Data were visualized in PRISM 10.

### Transcriptomic and Cytokine Assays

Isolated T cells were stimulated with anti-CD3/CD28 beads (500 nM, StemCell) and incubated in RPMI or ImmunoCult™ at 37°C for up to 48 hours. Media was removed from the cells and stored until analysis at -80°C. Cytokine secretion (TNF-α, IFN-γ, IL-4) was measured by multiplex bead-based ELISA using a ProcartaPlex™ Human Inflammation Panel (Thermo). The remaining cells were lysed for RNA extraction using RLT plus (Qiagen) and RNA from ELRi-isolated cells was extracted (RNeasy Plus Micro Kit, Qiagen). For RBC depletion experiments, the EasySep™ RBC Depletion Kit (StemCell) was used (Fig. 6).

### Transcriptomic Data Analysis

Bulk RNA sequencing was performed on red blood cells isolated from whole blood samples collected from 8 donors. For each donor, bench-based isolation method and the ELRi isolation method was applied with 16 samples total, with 8 replicates for each technique. Raw FASTQ files were generated through the Illumina Novaseq X sequencing platform. Read quality was assessed using fastQC and adapter trimming was performed using fastp (v101) (25). For our deconvolution analysis, trimmed reads were aligned and quantified using Kallisto (v0.51.1) with an index consisting of the human reference genome (GRCh38) and the genomes of common microbial contaminants, producing estimated sequencing pseudocounts. Microbial read fraction was evaluated to assess sample contamination. Post-quantification, human-aligned reads were categorized by transcript type (protein-coding, non-coding, and rRNA) based on GENCODE annotations. Sample quality was verified by assessing the burden of microbial reads and the proportion of non-protein-coding, mitochondrial, and ribosomal RNA transcripts. All samples met these quality control criteria and were included in the final analysis. The pseudocount matrix was filtered to remove transcripts that did not exhibit a read count of 5 or higher in at least 4 samples.

Differential gene expression (DGE) analysis was performed using DESeq2 (v1.44) (26) with follow-up log-2-fold shrinkage using the ashr package (v2.2.63) (27) in R 4.4.3. The bench isolation method was selected as reference. Log Genes with an absolute log2FC greater than 1 and an adjusted false discovery rate (FDR) < 0.05 were identified as differentially expressed. FDR was calculated using Benjamini-Hochberg test. A total of 28,626 genes were detected with 1,894 meeting the criteria as differentially expressed between the ELRi and bench isolation techniques. Heatmap was generated using Seaborn’s (v0.13.2) clustermap API for the top 25 differentially expressed genes ranked by their log2FCs, with unsupervised hierarchical clustering conducted on matrix of DE genes. Raw DESeq2 counts were log transformed through DESeq2 which were used for plotting the PCA and the heatmap, while log2FC was used to construct a volcano plot. Significantly differentially expressed genes, according to the prior criteria, are highlighted in red (upregulated in ELRi) or green (downregulated).

Cellular composition was estimated using AutoGeneS (v1.0.4) (28), fixed quantification deconvolution method). To accurately quantify Erythrocyte abundance, we compiled a combined reference set consisting of the Human PBMC single-cell data sourced from the Azimuth HubMAP Consortium (29), erythrocytes removed, and an erythrocyte reference consisting of specialized erythrocyte signatures derived from the red blood cell reference described in Jain et. al. 2022 (30). Imputed counts were visualized with stacked bar charts to show overall distributions and box-and-whisker plots to show per-cell-type differences in abundance. Mann-Whitney-U tests were used to identify statistically significant differences in imputed abundances by cell type using the statannotations (v0.7.2) package (31).

## Results

### ELRi Enables Efficient RBC Depletion and WBC Enrichment from Tiny Samples

Most standard immune cell isolation systems rely on milliliter volumes of blood, reducing their applicability for low volume samples. Blood is a complex fluid with white blood cells outnumbered 2000:1 by red blood cells (Figure 1A). While large scale purification steps usually rely on density gradient isolation to remove contaminating red blood cells, these procedures are not practical for microscale volumes. To perform an effective immune cell isolation from whole blood volumes as low as 8 μL, we developed a new process named exclusive liquid repellency isolation or ELRi. ELRi relies on the basic principles of exclusive liquid repellency (ELR) which demonstrated that an aqueous phase can be completely repelled from a solid phase when immersed in oil (19, 32, 33). Controlled by the balance of interfacial energies between solid, oil and liquid, ELR occurs when the sum of the interfacial energies of the solid/oil and the aqueous/oil interface energy results in a contact angle of <180° thus preventing the adhesion of the aqueous liquid to the surface (Figure 1B). Therefore, in ELRi, a stable aqueous droplet of blood is surrounded by silicone oil in a polypropylene tube (Figure 1C, D) which effectively prevents droplet adhesion and reduces RBC contamination. To isolate a specific immune cell type, the addition of antibodies and magnetic beads from commercial standard negative selection immune cell isolation kits followed by a brief incubation step then allows removal of unwanted red blood cells and immune components by using a custom-made microscale magnetic tip. The tip was constructed by embedding a 1/16” magnet into a standard P200 pipette tip. Upon contact of the magnetic tip with the droplet, magnetically labeled cells rapidly cluster toward the tip within approximately 1 minute, while unlabeled target cells remain at the bottom of the droplet. The removal of the magnetic tip removes the unwanted cells leaving only the purified immune cells remaining in the droplet. Details of the physics underlying the droplet splitting are provided in supplementary materials. Initial RBC:WBC ratios of ∼2000:1 were managed efficiently, producing a clarified cell suspension enriched in the desired immune subset. ELRi is compatible with volumes as low as 8 μL as shown in Figure 1C.

**Figure 1.**
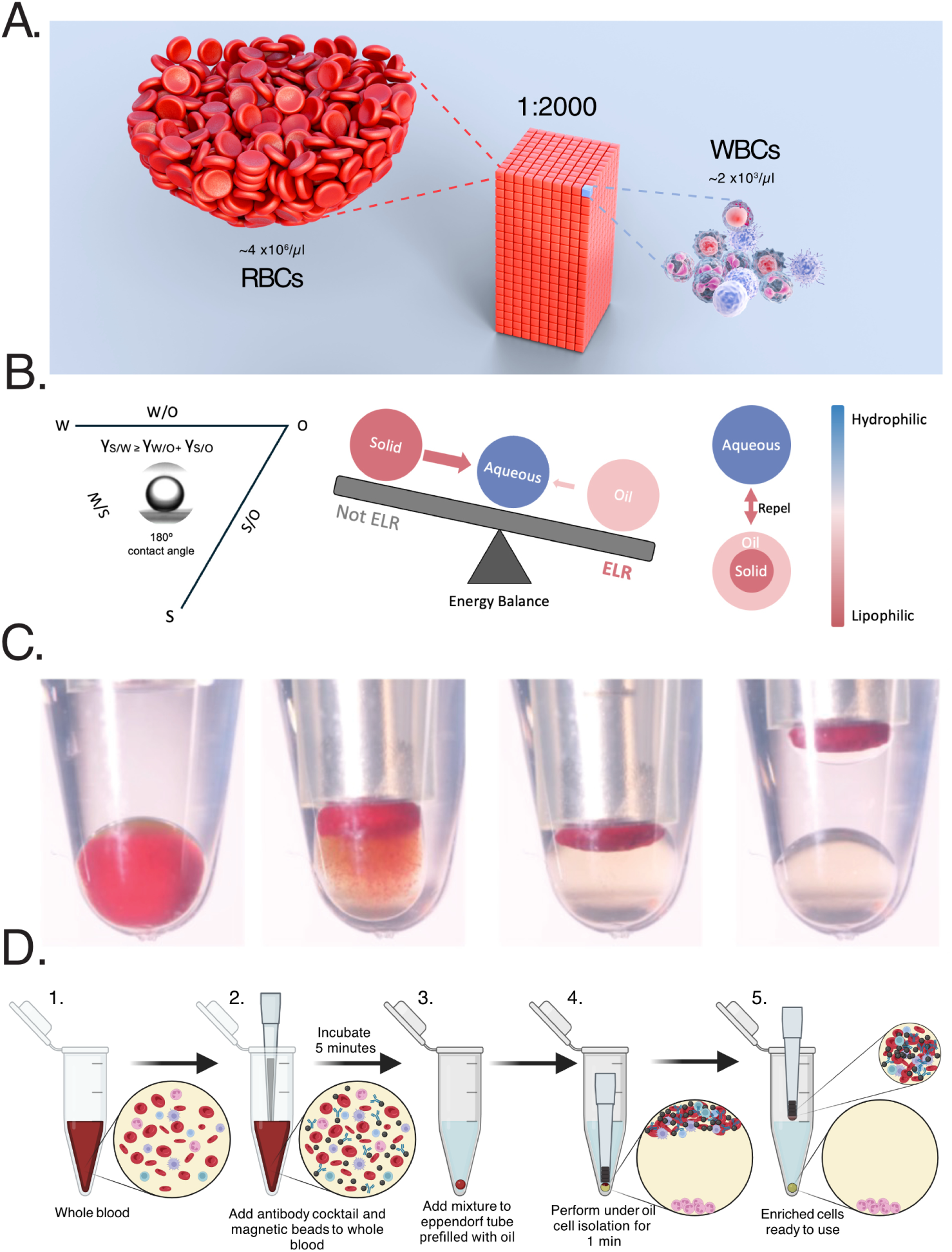
Implementation and principles of Exclusive Liquid Repellency-based isolation (ELRi). (a) Illustration of the typical cellular abundance disparity in whole blood (∼4×10^6^ RBCs/μL vs. ∼2×10^3^ WBCs/μL). This extreme imbalance usually necessitates complex microfluidics or large-scale magnetic isolation, highlighting ELRi’s potential as a simpler, more scalable alternative. (b) Conceptual illustration of the interfacial energy balance underlying exclusive liquid repellency (ELR). By considering the three-phase system, solid (S), aqueous (W), and oil (O), it is possible to delineate conditions where the dispersed aqueous phase completely repels the solid (i.e., a 180° contact angle), thus preventing adhesion. Achieving ELR ensures that the target cells remain in discrete aqueous droplets suspended at the oil interface rather than fouling the vessel surface. (c) Photographs demonstrating ELR-based droplet formation (8 μL blood) and enrichment of red blood cells, enabling easy droplet manipulation without fouling. (d) Schematic of the ELRi workflow: After adding a magnetic bead–antibody cocktail to whole blood, the mixture is introduced into a tube containing oil. Upon negative magnetic isolation, target cells are enriched at the droplet below, while non-target cells are removed.

### Precise control of ELRi using a Custom Magnet Rig

To enable precise control and facilitate reproducibility of ELRi, we designed and constructed a custom-built magnet rig. The rig is built around a Z axis rack and pinion stage that was modified to add a holder for the ELRi tube and custom-built magnet (Figure 2A, B). The rack and pinion mechanism enabled fine adjustments of the vertical position of the magnet in relation to the sample to provide control of the most effective distance for performing ELRi. The rig also standardizes these settings for specific volumes providing reproducible isolations. To further increase the capacity of ELRi, we manufactured disposable polypropylene sheaths (Figure 2C) that minimizes cross-contamination between samples and has the potential to allow multiple isolation rounds.

**Figure 2.**
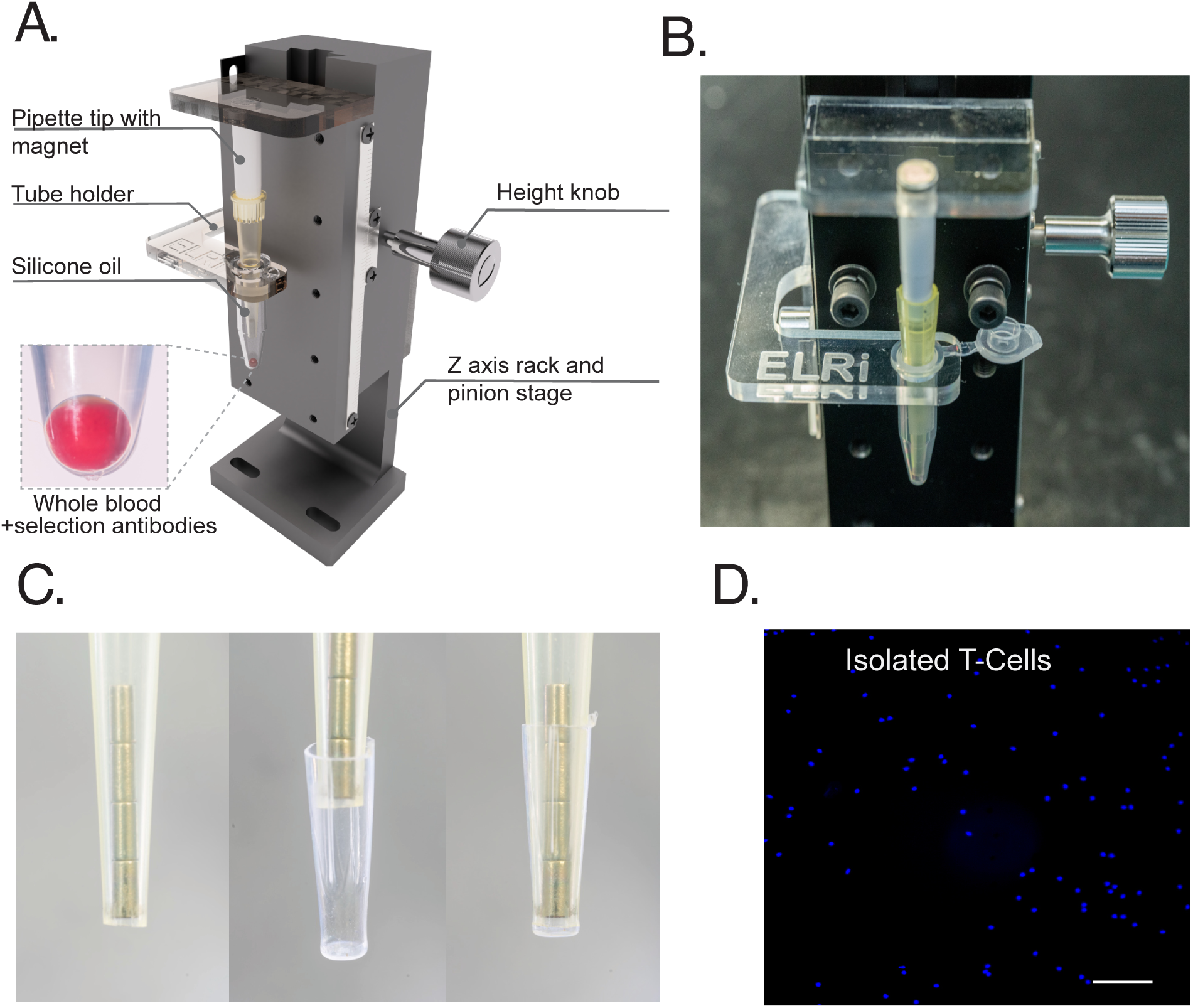
ELRi Rig and Workflow for Negative Isolation of Immune Cells with High Purity and Yield. (a) An ELRi isolation rig designed to perform isolation of cells within a standard 0.6 mL Eppendorf tube. The setup includes a fixed magnet and adjustable stage, allowing fine adjustments of the magnet to perform the isolation vertical. The rack-and-pinion mechanism enables fine adjustment of the magnet’s vertical position to optimize the isolation height. (b) Close-up view showing the rig and the Eppendorf tube secured to the rig. (c) A disposable, ELR-compatible polypropylene sheath prevents contamination between runs and allows rapid, selective enrichment of different target cell populations from the same initial sample. (d) Post-isolation Hoechst-stained images of purified T cells.

### ELRi-Isolation of T Cells

T cells play a central role in driving allergic disease inflammation, and the inception of disease via allergen-specific Th2 differentiation often occurs during early infancy. The underpinnings of this early-life allergen reactivity have been understudied due to the challenges of obtaining sufficient T cell numbers in very young children. Therefore, we initially chose to test ELRi for microscale isolation of T cells (Figure 2D). Further to confirm that ELRi isolated cells were of comparable purity, viability and functionality, we compared ELRi against a benchmark standard isolation method. Starting with a 10 mL venipuncture blood draw from healthy adult subjects, 30 μL of blood was removed and isolated using ELRi and a negative selection based magnetic T cell isolation kit while the remaining 10 mL was isolated using the same kit following the manufacturer’s standard recommendation. The isolated T cells were analyzed by flow cytometry to assess the purity (CD3) and viability (ghost red). ELRi isolated T cells with approximately 99% purity, were comparable to the benchmark isolation method, and viability was also equivalent between the two methods (Figure 3A). ELRi also reduced the isolation time to approximately 5 minutes, compared with approximately 35 minutes for the benchmark workflow. To assess whether ELRi-isolated T cells retained functional responses for downstream assays, T cells were isolated from the same blood samples from nine healthy adult donors using both ELRi and benchmark methods. To maintain comparable downstream culture conditions, isolated cell suspensions from both methods were volume matched by 1:1 dilution during isolation and cultured in the same 384-well plate format. Cells were cultured overnight with CD3/CD28 stimulation or left unstimulated as controls. Culture media was collected for cytokine analysis using multiplex bead-based ELISA, while cells were lysed for RNA extraction and qPCR analysis of Th1/Th2-associated cytokine genes, including IFN-γ, IL-4, and IL-13. CD3/CD28 stimulation increased cytokine-associated gene expression in both ELRi- and benchmark-isolated T cells, with comparable expression patterns between the two isolation methods (Figure 3B). Cytokine secretion analysis further showed substantial donor-to-donor variability in TNF-α, IFN-γ, and IL-4 production, but individual donor response patterns were generally consistent between ELRi and benchmark isolation methods (Figure 3C). Together, these data demonstrate that ELRi can isolate T cells from microliter-scale blood volumes with high purity and viability while preserving functional molecular and cytokine secretion responses comparable to standard macroscale purification methods.

**Figure 3.**
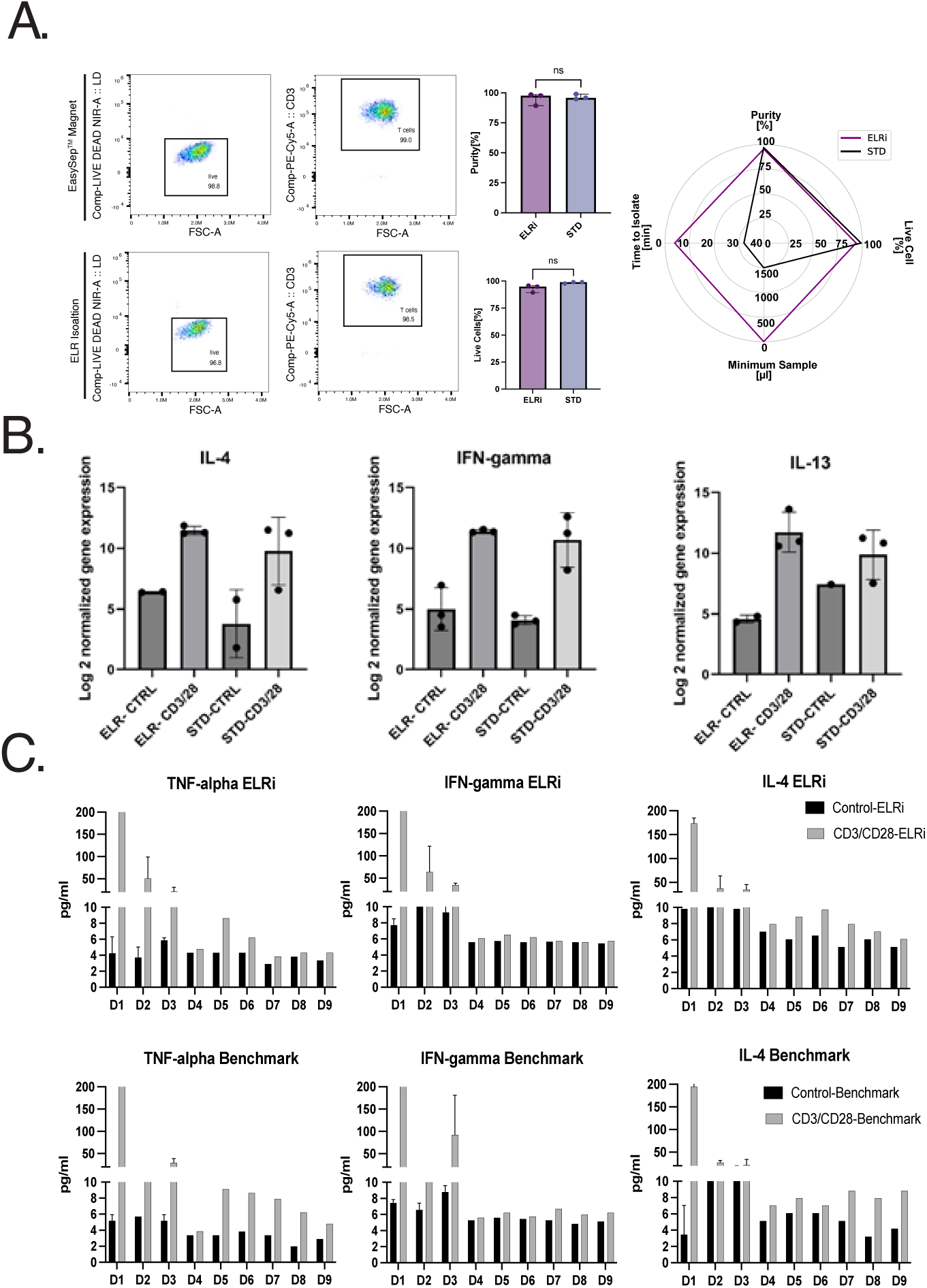
Cytokine Profiles, Gene Expression, and Purity Assessment of ELRi-Isolated T Cells. (a) Purity and viability assessment of ELRi-isolated T cells using flow cytometry. Radar plot shows reduced time and blood volume required for ELRi compared to standard methods. (b) Gene expression analysis of IL-4, IFN-γ, and IL-13 using qPCR. ELRi-isolated T cells maintain consistent gene expression levels compared to standard isolation methods, supporting the effectiveness of ELRi in preserving cell functionality. (c) Cytokine secretion profiles for TNF-α, IFN-γ, and IL-4 from nine different donors in ELRi-isolated (top) versus benchmark-isolated (bottom) T cells. ELRi-isolated T cells (gray bars) stimulated with CD3/CD28 exhibit comparable or improved cytokine production compared to controls (black bars), demonstrating functional integrity post-isolation.

### Scaling ELRi to Accommodate Larger Droplet Volumes (ELRi-XL)

To further expand the applicability of ELRi beyond lymphocyte populations, we investigated whether the platform could be scaled to enable isolation of rarer cell types from small volumes of blood. While T cells comprise the majority of lymphocytes in whole blood (∼70%) and served as the initial population characterized using ELRi, isolation of rarer populations presents additional technical constraints due to limited starting material. To address this limitation, we developed an expanded format of the platform, termed ELRi-XL, designed to accommodate larger droplet volumes (∼30 μL and beyond). ELRi-XL employs a 5 mL Axygen tube in combination with 200 μL of silicone oil (Figure 4A, B). The magnet rig was modified to securely hold the larger tube, and the optimal magnet–droplet distance for efficient isolation was empirically determined.

**Figure 4.**
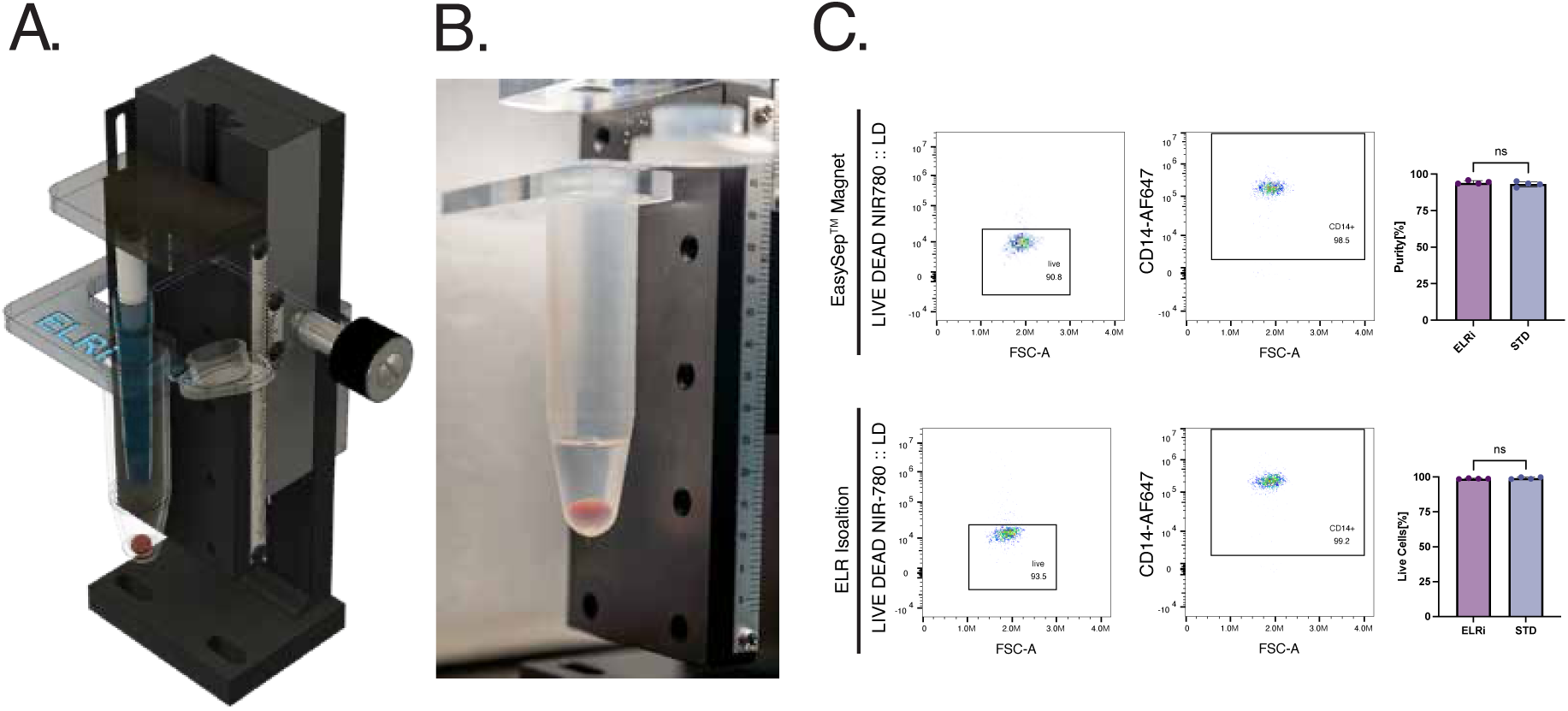
Scaling Up ELRi for Isolation of less prevalent Cell Types. (a) Schematic of the scaled-up ELRi rig configured for processing larger sample volumes to isolate rare cell types such as monocytes. The setup utilizes a 5 mL Axygen tube in place of the standard 0.6 mL Eppendorf tube, with a scaled-up magnet housed within a 1000 µL pipette tip for enhanced magnetic separation efficiency. (b) Photograph of the scaled-up setup in operation. The larger Axygen tube accommodates 70 µL droplets of whole blood per assay, enabling the isolation of rarer cell populations while maintaining a streamlined workflow. (c) Representative flow cytometry analysis demonstrating the performance of scaled-up ELRi isolation compared to standard magnetic separation (“EasySep™ Magnet”) for isolating rare cell populations. Dot plots depict gating strategies for live cells identified via LIVE/DEAD NIR780 viability staining (left plots). Middle plots show forward scatter (FSC-A) versus CD14-AF647 staining for monocyte identification, with gates illustrating the percentage of CD14-positive monocytes obtained using each method (EasySep™, top; ELRi, bottom). Purity percentages are indicated within gates. Right bar graphs quantitatively summarize monocyte purity and cell viability, confirming no significant differences (ns, p > 0.05) between scaled-up ELRi and standard magnetic isolation (STD). These results validate that the scaled-up ELRi maintains comparable purity and viability for rare cell isolation, suitable for subsequent downstream functional assays.

To validate ELRi-XL, we isolated monocytes, a less prevalent leukocyte population (∼20% of peripheral blood mononuclear cells), from a 30 μL droplet of whole blood using a commercial monocyte isolation kit. Flow cytometry analysis demonstrated high purity (99.2%) and viability (93.5%) of recovered monocytes. Although differences were not statistically significant, ELRi-XL yielded slightly higher purity (99.2% vs 98.6%) and viability (93.5% vs 90.8%) compared to a conventional benchmark isolation performed on 10 mL of blood using the same kit (Figure 4C). These results demonstrate that ELRi can be scaled to match blood volume availability and experimental needs without compromising isolation efficiency or cell quality.

### Microscale Functional Analysis of Pediatric T Cells Using ELRi

Having established the scalability of ELRi, we next evaluated its ability to support downstream functional assays using clinically relevant pediatric samples. We designed a study to assess the function and phenotype of T cells in children with asthma.

Blood samples were collected from children aged 4–16 years undergoing a routine blood draw for clinical reasons at the outpatient clinic, with 11 children in allergic, non-asthmatic group and 10 in the allergic, asthmatic group (Table 1). A 30 μL volume of whole blood was used for T-cell isolation via ELRi. Isolated T cells were recovered from the ELRi droplet for quantification of migratory responses.

**Table 1.** Subject Characteristics.

| n | 10 | 11 |
| --- | --- | --- |
| History of allergy | Yes | Yes |
| History of asthma | No | Yes |
| **Demographics** |  |  |
| Age (yrs) | 7-16 | 4-12 |
| Gender Male n (%) | 7 (70%) | 9 (82%) |
| Gender Female n (%) | 3 (30%) | 2 (18%) |
| **Race/ Ethnicity** |  |  |
| Hispanic | 0 (0%) | 1 (9%) |
| Mixed or other | 3 (30%) | 2 (18%) |
| White | 7 (70%) | 8 (73%) |

The ELRi isolated T cells were immediately introduced into channels of the KOALA platform pre-coated with ICAM-1 (Figure 5A, B). A KOALA lid containing collagen mixed with either CCL21 or PBS as a control was then applied. Time-lapse imaging was performed over 1.5 hours, and T cell migration was quantified using a custom-built image analysis pipeline to identify and track individual cells across the field of view (Figure 5C).

**Figure 5.**
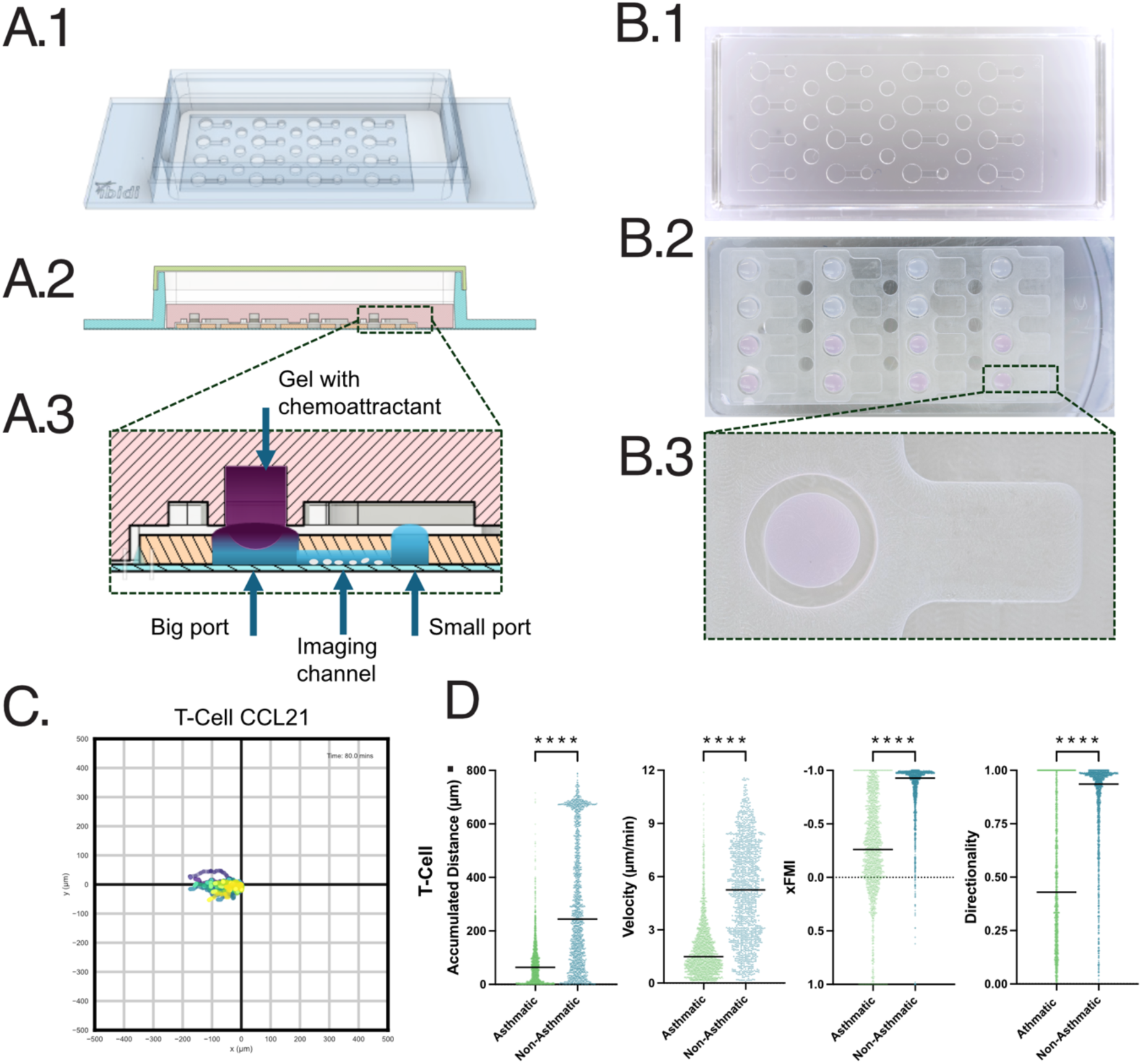
Migration Analysis Using the KOALA Platform. (a) Schematic representation of the KOALA (Kit-On-A-Lid Assay) platform for cell migration studies. (a.1) The assembled KOALA device designed for cell migration analysis. (a.2) Cross-sectional view of the platform showing the layout of the wells. (a.3) Detailed close-up of the assay chamber (b.1) Overview of the device showing multiple wells for parallel assays. (b.2) Magnified view of the wells loaded with samples. (b.3) Close-up of a single well. (c) Polar plots of T cell migration in response to chemotactic gradients. (d) Quantitative analysis of migration using the forward migration index (xFMI)) metric, velocity, distance and directionality. Plots show significant increase in xFMI, migration distance, and velocity between non-asthmatic and asthmatic T cells. Notably, the directionality in the asthmatic donor group is impaired, highlighting a potential deficit in chemotactic response in this cohort.

T cells exhibited directional migration toward the CCL21 chemoattractant (Figure 5D). Quantitative analysis of the Foward Migration Index (FMI) representing the mean endpoint of all tracked cells, revealed that T cells from non-asthmatic children migrated significantly farther toward the chemokine stimulus than those from asthmatic children (Figure 5D). Migration distance and velocity were also significantly impaired in the asthmatic children compared to the non-asthmatic children. Interestingly, while T cells from non-asthmatic children migrated strongly toward the CCL21 stimulus, the T cells from asthmatic children showed less directionality towards the stimulus (Figure 5D). Collectively, these data demonstrate that ELRi enables isolation of functional T cells from very small volumes of pediatric blood and supports downstream microscale functional assays relevant to disease phenotyping.

### Enhanced Transcriptomic Resolution Post-RBC Depletion

A persistent limitation of whole-blood RNA-seq is that erythrocyte- and globin-derived transcripts can account for a disproportionate share of reads, reducing effective library complexity for leukocyte transcripts. To test if ELRi could deplete red blood cells from microliter volume blood samples, we collected 30 μL blood samples and either directly lysed cells for RNA extraction (benchmark control) or isolated white cells by ELRi followed by lysis for RNA (ELRi). This is directly reflected in the top-DEG heatmap (Fig. 6A), where benchmark libraries are dominated by erythrocyte/haemoglobin-associated genes (boxed). In contrast, ELRi-treated libraries show consistent absence of these erythroid features across donors (Figure 6A), indicating that the primary effect of ELRi is depletion of high-abundance RBC transcript signal.

**Figure 6.**
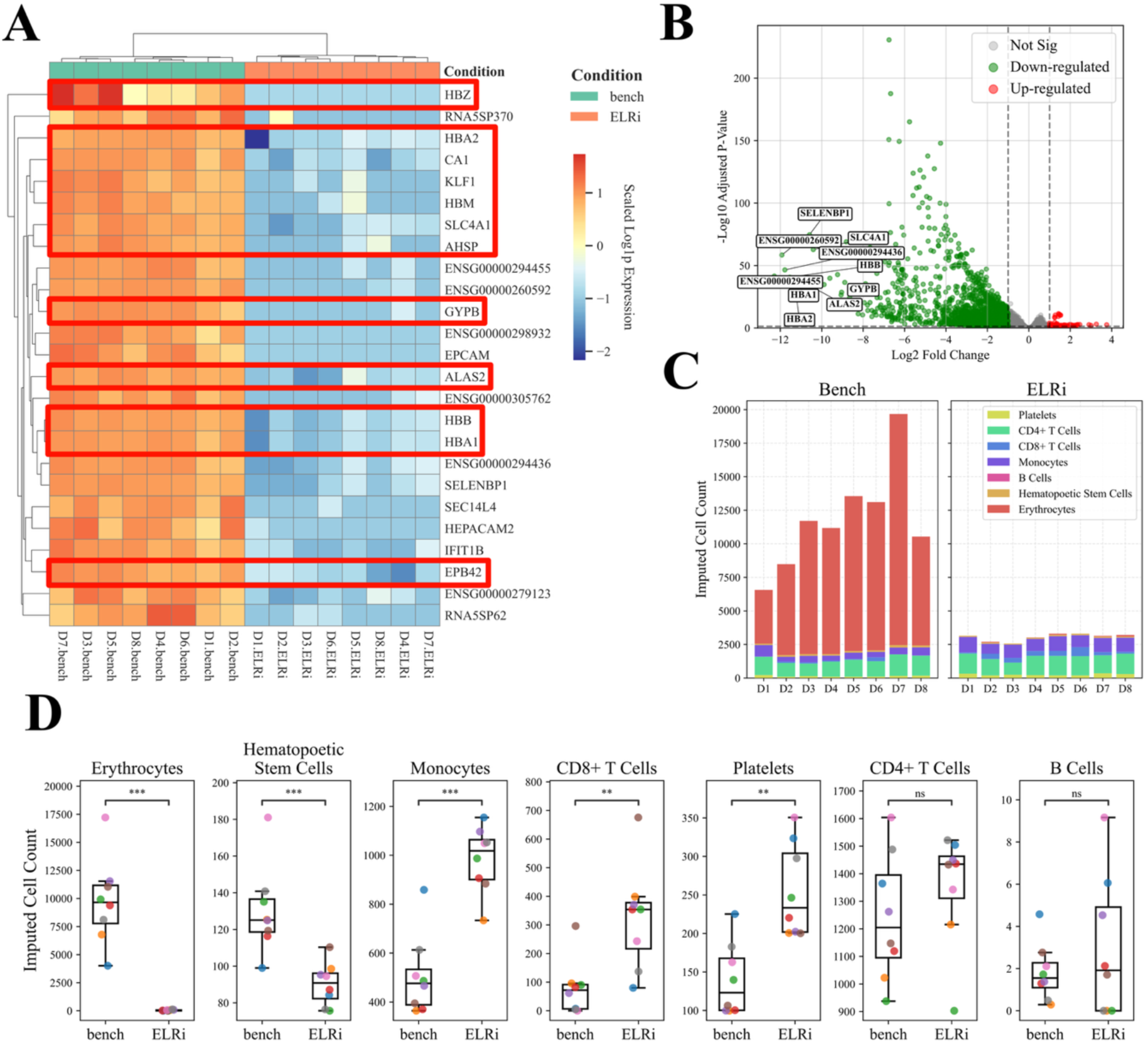
ELRi globin-depletion improves RNA-seq library complexity and sensitivity. A) Heat-map of the 25 most significant DEGs (variance-stabilised counts, row z-scores). Bench libraries (teal) are dominated by erythrocyte/haemoglobin genes (boxed), whereas ELRi (salmon) markedly suppresses these reads. B) Volcano plot (DESeq2, |log₂FC| > 1, FDR < 0.05). Significant genes are almost exclusively downregulated under ELRi and correspond to globin/RBC transcripts. C) Cell-type composition estimates from bulk RNA-seq deconvolution, showing a strong reduction in the erythrocyte component in ELRi-treated samples with relative enrichment of immune-relevant populations (e.g., CD4 T, CD8 T, and monocytes). D) Box plots of cell-type deconvolution for the top 7 most abundant populations. P-values calculated via Mann-Whitney U test (***p < .001, **p < .01). ELRi samples show significantly lower abundance of erythrocytes and HSCs, but increased levels of monocytes, CD8+ T cells, and platelets.

This depletion results in a clear exposure-driven separation of expression profiles. Differential expression results further support the specificity of the change: the volcano plot shows that the most significant changes under ELRi are predominantly downregulated features (Figure 6B), aligning with absence of erythrocyte/globin-associated transcripts rather than widespread bidirectional shifts. Orthogonal composition analyses provide additional context for interpretation. Deconvolution-based estimates show a marked reduction of the erythrocyte component in ELRi libraries and a relative increase in immune-relevant populations including platelets, CD8 T cells, and monocytes (Figure 6C). Consistently, enrichment-style summaries show diminished erythrocyte identity signal and clearer immune-associated patterns in ELRi samples across donors (Figure 6D). Taken together, (panels A-D) these data demonstrate ELRi reduces RBC-derived transcript dominance, which allows for leukocyte transcriptional analyses that are typically of primary interest in immunology and clinical profiling. Further, resolution of platelet numbers was also increased, which could support low volume platelet-sequencing applications for biomarkers in oncology or inflammation.

This is particularly relevant for ultra-low input and volume-limited studies, where RBC transcript depletion is not routinely incorporated as a standard preprocessing step, and investigators often rely on deeper sequencing or post hoc normalization to contend with globin dominance. In contrast, the results here show that an upstream depletion step can substantially reduce erythroid signal (Figure 6A, C) and its contribution to sample variance (Figure 6B), while improving the relative detectability of immune-associated programs (Figure 6D). For studies constrained by limited blood volume or limited recoverable RNA, such as pediatric sampling, longitudinal monitoring, or scarce biobanked aliquots, this approach offers a practical means to improve interpretability without presuming access to larger inputs or substantially higher sequencing depth.

## Discussion

Here we report the ELRi platform as a new tool for isolation of immune cells from microscale blood samples. ELR describes the condition in which an aqueous droplet is completely repelled (contact angle ≈ 180°) from a solid surface by an immiscible continuous phase (often oil) (19). In a three-phase system consisting of a solid (S), a water-based liquid (W), and an oil (O), this behavior is governed by the interfacial energies (γ) between each pair of phases. Specifically, when the following inequality is satisfied, γS/W + γW/O ≤ γS/O, the aqueous droplet is unable to wet the solid, instead remaining fully suspended within the oil phase. To create a straightforward system that does not require specialized equipment we leveraged the properties of standard polypropylene tubes which have inherent ELR when combined with an appropriate oil overlay. This configuration creates a droplet of aqueous sample that never contacts the container walls, preventing evaporation and sample loss. In volumes as low as 8 μL, this stable droplet environment supports direct magnetic bead-based isolation of immune cells without the complexities of specialized microfluidic chips or large initial volumes. Thus, by creating a stable aqueous droplet suspended in silicone oil, ELRi leverages exclusive liquid repellency to prevent sample loss, biofouling and contamination. This innovation allowed us to efficiently remove RBCs and isolate blood cell subsets, even from just a few microliters of blood, without sacrificing cellular health or functionality.

In early life, stimulation of immune pathways through environmental factors can drive Th2-biased polarization and subsequent allergy development (34). While several risk factors have been identified, acquisition of blood samples from neonates in sufficient volumes to permit functional immune assays and molecular characterization is challenging. Using the combination of ELRi and KOALA, here we demonstrated that we could perform functional analysis of T cells from 30 μL of blood. KOALA consists of a microscale straight channel base with a volume of ∼150nl, and a custom-built lid filled with collagen droplets containing reagents for the assay i.e. chemoattractants (12). Immune cells can be loaded into the channel and when paired with the lid, fluid is transferred between the lid and base beginning the functional assay. KOALA has been used to show robust measurement of neutrophil migration (12), with neutrophils directly captured in the device. However, this analysis has not been applied to less prevalent immune subsets such as T cells.

ELRi demonstrated that the isolated T cell function remained intact, as shown by their robust cytokine secretion and consistent gene expression patterns when compared to the benchmark samples (Figure 3). Additionally, integrating ELRi-isolated cells into the KOALA platform (Figure 5) demonstrated preserved chemotactic responses, suggesting that sensitive cellular behaviors are unaffected by the isolation process. This was particularly insightful when analyzing cells from pediatric asthmatic donors, where T cells exhibited fundamentally suppressed migratory behavior compared to non-asthmatic controls. Further, T cells from asthmatic individuals lacked directionality of migration towards the CCL21 chemokine stimulus, suggesting a defect in CCR7-mediated trafficking that could impair immune surveillance and contribute to chronic inflammation or impaired responses to respiratory viruses (35). These novel findings suggest circulating T cells in asthmatic children have impaired CCR7 receptor function or altered CCR7 expression. CCR7 is expressed on naïve and central memory T cells. While quantification of CCR7 expression was beyond the scope of this study, with age-matched pediatric participants significant differences in naïve and central memory T cell frequencies would not be expected.

Some limitations should be considered for these findings. The sample size in the asthmatic and non-asthmatic groups were relatively small and taken from older children. Additional work should focus on analysis of a larger group of patients. Measurements of CCL21 in serum would also be a valuable addition.

The flexibility of ELRi is a major advantage as it can be applied to any immune cell types that have magnetic bead-based isolation kits available and the scaling up droplet size enabled the retrieval of less frequent subsets such as monocytes (Figure 4), expanding the platform’s applicability. Depletion of RBCs is another advantage of the platform. While standard input cDNA library kits are compatible with globin removal, these methods are not applicable to ultra-low input kits needed for microliter volume samples. We demonstrated that RBCs could be effectively and quickly removed from blood samples prior to lysis for RNA extraction and sequencing. Globin genes were significantly reduced and the immune cell transcripts enriched in the ELRi samples (Figure 6) demonstrating that with this platform, 30 μL is sufficient to return good quality transcriptomic data.

Compared to conventional isolation techniques that require milliliter-scale samples, time-consuming steps, or complex instrumentation, ELRi offers a rapid, accessible, and versatile alternative. The approach is well suited for pediatric, neonatal, and other patient populations where sample sparing is critical. By preserving cell viability and providing immediate downstream functional assessment compatibility, ELRi supports a wide range of analyses, from immune profiling and migration assays to transcriptomics and functional screening, in a fraction of the volume typically required. In conclusion, ELRi represents an enabling tool for the microliter isolation and functional characterization of immune cells. Its simplicity, scalability, and compatibility with advanced assay platforms like KOALA open new avenues for high-frequency, low-burden sampling. By facilitating the discovery of functional alterations, such as the chemotactic deficiencies observed in asthmatic immune cells, the platform allows for more precise and individualized immunological assessments.

## Supporting information

Details of the physics underlying the droplet splitting are provided in supplementary materials

## Acknowledgements

This work was supported by NIH U24AI152177 and the University of Wisconsin Carbone Cancer Center Support Grant NIH P30CA014520. We thank the University of Wisconsin Carbone Cancer Center Flow Cytometry Laboratory, supported by P30 CA014520, for use of its facilities and services.

## Conflict of interest disclosure

David J. Beebe holds equity in Bellbrook Labs LLC, Salus Discovery LLC, Lynx Biosciences Inc., Stacks to the Future LLC, Flambeau Diagnostics LLC, Eolas Diagnostics, Inc., Navitro Biosciences LLC, and Onexio Biosystems LLC. The other authors have no conflicts of interest to report.

## Notes

### Summary of Updates

The previous version had a problem with figure legibility. This version corrects this problem.

## References

1. Debes GF, Arnold CN, Young AJ, Krautwald S, Lipp M, Hay JB, et al. Chemokine receptor CCR7 required for T lymphocyte exit from peripheral tissues. Nature immunology. 2005;6(9):889–94.

2. Syed F, Blakemore SJ, Wallace DM, Trower MK, Johnson M, Markham AF, et al. CCR7 (EBI1) receptor down-regulation in asthma: differential gene expression in human CD4+ T lymphocytes. QJM. 1999;92(8):463–71.

3. El Brihi J, Pathak S. Normal and Abnormal Complete Blood Count With Differential. StatPearls. Treasure Island (FL)2026.

4. Nivedita N, Papautsky I. Continuous separation of blood cells in spiral microfluidic devices. Biomicrofluidics. 2013;7(5):54101.

5. Choi J, Hyun JC, Yang S. On-chip Extraction of Intracellular Molecules in White Blood Cells from Whole Blood. Sci Rep. 2015;5:15167.

6. Wu L, Guan G, Hou HW, Bhagat AA, Han J. Separation of leukocytes from blood using spiral channel with trapezoid cross-section. Anal Chem. 2012;84(21):9324–31.

7. Agrawal N, Toner M, Irimia D. Neutrophil migration assay from a drop of blood. Lab Chip. 2008;8(12):2054–61.

8. Berthier E, Guckenberger DJ, Cavnar P, Huttenlocher A, Keller NP, Beebe DJ. Kit-On-A-Lid-Assays for accessible self-contained cell assays. Lab Chip. 2013;13(3):424–31.

9. Ellett F, Jorgensen J, Marand AL, Liu YM, Martinez MM, Sein V, et al. Diagnosis of sepsis from a drop of blood by measurement of spontaneous neutrophil motility in a microfluidic assay. Nat Biomed Eng. 2018;2(4):207–14.

10. Moussavi-Harami SF, Mladinich KM, Sackmann EK, Shelef MA, Starnes TW, Guckenberger DJ, et al. Microfluidic device for simultaneous analysis of neutrophil extracellular traps and production of reactive oxygen species. Integr Biol (Camb). 2016;8(2):243–52.

11. Otawara M, Roushan M, Wang X, Ellett F, Yu YM, Irimia D. Microfluidic Assay Measures Increased Neutrophil Extracellular Traps Circulating in Blood after Burn Injuries. Sci Rep. 2018;8(1):16983.

12. Sackmann EK, Berthier E, Young EW, Shelef MA, Wernimont SA, Huttenlocher A, et al. Microfluidic kit-on-a-lid: a versatile platform for neutrophil chemotaxis assays. Blood. 2012;120(14):e45–53.

13. Wu J, Hillier C, Komenda P, Lobato de Faria R, Levin D, Zhang M, et al. A Microfluidic Platform for Evaluating Neutrophil Chemotaxis Induced by Sputum from COPD Patients. PLoS One. 2015;10(5):e0126523.

14. Elsemary MT, Maritz MF, Smith LE, Warkiani ME, Thierry B. Two-stage inertial microfluidics enrichment of activated T-cells towards a bead-less chimeric antigen receptor manufacturing protocol. Med Oncol. 2026;43(3):126.

15. Howie SR. Blood sample volumes in child health research: review of safe limits. Bull World Health Organ. 2011;89(1):46–53.

16. Cannon CA, Ramchandani MS, Golden MR. Feasibility of a novel self-collection method for blood samples and its acceptability for future home-based PrEP monitoring. BMC Infect Dis. 2022;22(1):459.

17. Hendelman T, Chaudhary A, LeClair AC, van Leuven K, Chee J, Fink SL, et al. Self-collection of capillary blood using Tasso-SST devices for Anti-SARS-CoV-2 IgG antibody testing. PLoS One. 2021;16(9):e0255841.

18. Wickremsinhe E, Fantana A, Berthier E, Quist BA, Lopez de Castilla D, Fix C, et al. Standard Venipuncture vs a Capillary Blood Collection Device for the Prospective Determination of Abnormal Liver Chemistry. J Appl Lab Med. 2023;8(3):535–50.

19. Li C, Yu J, Schehr J, Berry SM, Leal TA, Lang JM, et al. Exclusive Liquid Repellency: An Open Multi-Liquid-Phase Technology for Rare Cell Culture and Single-Cell Processing. ACS Appl Mater Interfaces. 2018;10(20):17065–70.

20. Jang JS, Berg B, Holicky E, Eckloff B, Mutawe M, Carrasquillo MM, et al. Comparative evaluation for the globin gene depletion methods for mRNA sequencing using the whole blood-derived total RNAs. BMC Genomics. 2020;21(1):890.

21. Shin H, Shannon CP, Fishbane N, Ruan J, Zhou M, Balshaw R, et al. Variation in RNA-Seq transcriptome profiles of peripheral whole blood from healthy individuals with and without globin depletion. PLoS One. 2014;9(3):e91041.

22. Harrington CA, Fei SS, Minnier J, Carbone L, Searles R, Davis BA, et al. RNA-Seq of human whole blood: Evaluation of globin RNA depletion on Ribo-Zero library method. Sci Rep. 2020;10(1):6271.

23. Sheerin D, Lakay F, Esmail H, Kinnear C, Sansom B, Glanzmann B, et al. Identification and control for the effects of bioinformatic globin depletion on human RNA-seq differential expression analysis. Sci Rep. 2023;13(1):1859.

24. Juang TD, Riendeau J, Geiger PG, Datta R, Lares M, Yada RC, et al. Micro blood analysis technology (μBAT): multiplexed analysis of neutrophil phenotype and function from microliter whole blood samples. Lab on a Chip. 2024;24(17):4198–210.

25. Chen S, Zhou Y, Chen Y, Gu J. fastp: an ultra-fast all-in-one FASTQ preprocessor. Bioinformatics. 2018;34(17):i884–i90.

26. Love MI, Huber W, Anders S. Moderated estimation of fold change and dispersion for RNA-seq data with DESeq2. Genome Biol. 2014;15(12):550.

27. Stephens M. False discovery rates: a new deal. Biostatistics. 2017;18(2):275–94.

28. Aliee H, Theis FJ. AutoGeneS: Automatic gene selection using multi-objective optimization for RNA-seq deconvolution. Cell Syst. 2021;12(7):706–15 e4.

29. Hao Y, Stuart T, Kowalski MH, Choudhary S, Hoffman P, Hartman A, et al. Dictionary learning for integrative, multimodal and scalable single-cell analysis. Nat Biotechnol. 2024;42(2):293–304.

30. Jain V, Yang WH, Wu J, Roback JD, Gregory SG, Chi JT. Single Cell RNA-Seq Analysis of Human Red Cells. Front Physiol. 2022;13:828700.

31. Charlier F, Weber M, Izak D, Harkin E, Magnus M, Lalli J, et al. Statannotations. v0.6 ed: Zenodo; 2022.

32. Li C, Humayun M, Walker GM, Park KY, Connors B, Feng J, et al. Under-Oil Autonomously Regulated Oxygen Microenvironments: A Goldilocks Principle-Based Approach for Microscale Cell Culture. Adv Sci (Weinh). 2022;9(10):e2104510.

33. Li C, Yu J, Paine P, Juang DS, Berry SM, Beebe DJ. Double-exclusive liquid repellency (double-ELR): an enabling technology for rare phenotype analysis. Lab Chip. 2018;18(18):2710–9.

34. Cereta AD, Oliveira VR, Costa IP, Guimaraes LL, Afonso JPR, Fonseca AL, et al. Early Life Microbial Exposure and Immunity Training Effects on Asthma Development and Progression. Front Med (Lausanne). 2021;8:662262.

35. Kloepfer KM, Jackson DJ, Jartti T, Liu AH, Gern JE. How Infections and Immune Development Relate to Preschool Recurrent Wheezing and Asthma. J Allergy Clin Immunol Pract. 2025;13(10):2553–61.

