## Supplementary material for "Exclusive liquid repellency isolation (ELRi) microliter scale isolation technique unmasks T-cell migration defects to CCL21 in pediatric asthma": Details of the physics underlying the droplet splitting are provided in supplementary materials

**Rationale for Deriving Droplet Splitting Physics**

Reproducible operation of ELRi requires that magnetically labeled blood sub-fractions (e.g., an RBC-rich portion) separate in a controlled and repeatable manner, without unintended fragmentation or bulk droplet displacement. In our current workflow, the target pinch-off volumes and operating conditions were initially established empirically; however, empirical tuning alone can be sensitive to operator technique, sample-to-sample variability (e.g., hematocrit and bead loading), and modest differences in magnet geometry or placement. A quantitative description of the splitting process is therefore essential to define the parameter space in which ELRi can be replicated reliably across experimental runs and users.

To this end, we derive a force-balance framework that (i) predicts the conditions under which the magnetic force applied at the droplet interface exceeds the capillary force required for pinch-off, and (ii) delineates the regimes in which the droplet remains gravitationally stable within the tube. These relationships provide design rules for selecting initial droplet volume and labeled fraction, and for specifying the required magnetic field–gradient product at the tip such that the pinch-off event occurs consistently. In addition, the framework enables systematic evaluation of how droplet size, whole-blood-to-diluent ratio (i.e., the fraction of the RBC-rich portion), and applied magnetic strength jointly influence the isolation outcome, thereby supporting rational optimization rather than trial-and-error adjustment. Collectively, this analysis also allows verification that the effective magnetic forcing at the tip is repeatable under standardized magnet placement and hardware configuration, improving inter-operator consistency and overall robustness of the ELRi phenomenon.

**Droplet Splitting Physics**

This section derives the force–balance criteria under which a magnetically labeled blood sub–droplet (e.g. RBC–rich) is pulled away (“pinched off”) from the main droplet, as well as the conditions under which the droplet remains stable at the bottom of the tube under gravity.

**Magnetic Pinch–Off Condition**

Let the total droplet have volume *V* , of which a fraction *f* (where 0 *< f <* 1) is occupied by the magnetically labeled RBC–rich portion. For instance, *f* = 1*/*2 might represent a 1:1 RBC–to– buffer ratio, whereas *f* = 1*/*4 could represent a 1:3 dilution. We approximate each portion (main droplet vs. RBC–rich sub–droplet) as nearly spherical, so the sub–droplet volume is *fV* and its characteristic radius is

*R*_RBC_
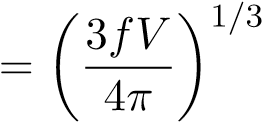
 *.*	(1)

To detach this RBC–laden sub–droplet, the *magnetic force* pulling on the cells must exceed the *capillary force* that holds the droplet together. A simple estimate for the capillary force required to “pinch off” a sub–droplet of radius *R*_RBC_ is

*F*cap ≈ 2*πR*RBC*γ,*	(2)

where *γ* is the effective blood/oil interfacial tension.

The magnetic force acting on the labeled RBC–rich portion can be written as

*F*_mag_ = *µ*_0_*χ*_eff_(*fV* )(**B** · ∇)**B***,*	(3)

where *µ*_0_ is the permeability of free space, *χ*_eff_ is an effective magnetic susceptibility capturing bead loading, and (**B** · ∇)**B** represents the local magnetic field–gradient product.

Pinch–off occurs when

*F*_mag_ *> F*_cap_	=⇒	*µ*_0_*χ*_eff_(*fV* )(**B** · ∇)**B** *>* 2*πR*_RBC_*γ.*	(4)

Substituting the expression for *R*_RBC_ yields a threshold condition on the magnetic field strength and gradient required to detach an RBC–rich fraction of volume *fV* . In practice, small NdFeB magnets (3–8mm diameter) placed a few millimeters from the droplet can generate fields of 0.3–

0.5T with gradients of 10–100Tm^−1^, often satisfying this criterion.

**Droplet Stability Under Gravity**

The silicone oil creates an ultrathin nonwetting layer between the blood droplet and the polypropylene tube, yielding an effective contact angle approaching 180^◦^ and negligible wall adhesion. Nevertheless, the droplet remains at the bottom of the tube due to gravity.

The relative importance of gravitational and interfacial forces is quantified by the Bond number,

Bo =
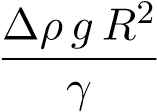
*,*	(5)

where ∆*ρ* is the density difference between blood and silicone oil, *g* is gravitational acceleration, *R* = (3*V/*4*π*)^1^*^/^*^3^ is a characteristic droplet radius, and *γ* is the interfacial tension.

For Bo ≪ 1, surface tension dominates and the droplet remains nearly spherical. Even for Bo ∼ 1, gravity only slightly flattens the droplet, and the large contact angle prevents wetting or wall climbing. Since blood is denser than 5cSt silicone oil, buoyancy does not drive upward motion. Thus, in the absence of external forces exceeding ∆*ρgV* or the capillary pinch–off force, the droplet remains stably seated at the bottom of the tube.
